# Evaluating novel sources of resistance to *Fusarium oxysporum* f. sp. *Cubense* tropical race 4 (TR4) in a global banana diversity panel

**DOI:** 10.64898/2026.11.18.767592

**Authors:** Carolina Olave-Achury, Daniel Ponndorf, Angela Chaparro Garcia, Anthony Hall, Graham McGrann, Jose J De Vega

## Abstract

*Fusarium oxysporum* f. sp. *cubense* tropical race 4 (Foc TR4) poses a major threat to global banana production, yet quantitative variation in resistance across *Musa* diversity remains poorly characterised. In this study, 21 accessions representing many of the major cultivated *Musa* groups and wild relatives were evaluated for their response to Foc TR4 under controlled-environment conditions. Significant variation in Foc TR4-associated corm discolouration was observed across the panel. Eight accessions had significantly less corm discolouration than the susceptible control Dwarf Cavendish: five were classified as resistant (all plants with more than 80% healthy corm area): *Musa itinerans*, *M. schizocarpa*, FHIA-02, FHIA-21 and the plantain Curare, and three as tolerant, which response was significantly different to the Dwarf Cavendish control but with some plants affected: *M. ornata*, Pelipita and Mai’a Maoli Koa’e. All accessions with only A genomes were susceptible or very susceptible, as were *M. textilis* and the B-containing triploids Pome (AAB) and Silver Bluggoe (ABB). Five of the seven accessions carrying at least one B genome were resistant or tolerant, suggesting that B-genome backgrounds merit wider screening as potential sources of resistance. These results identify valuable sources of resistance to Foc TR4 and provide a basis for banana breeding and further investigation of the genetic basis of resistance at the cultivar level.

## Introduction

Bananas, members of the genus *Musa*, are large herbaceous plants cultivated across tropical and subtropical regions of Southeast Asia, Africa, and the Americas. They are among the most important crops worldwide, with more than 183 million tonnes of bananas and plantains produced in 2023 (FAO 2025). The domestication and selection of seedless fruit led to the fixation of parthenocarpy and sterility in cultivated bananas. As a result, these crops are propagated mainly vegetatively. At the same time, thousands of years of diversification among wild genotypes, together with human selection of hybrids, have given rise to the wide range of edible bananas and plantains available today. More than 7,000 *Musa* accessions are preserved in global germplasm collections curating genetic diversity derived from geographical adaptation and local selection (Ruas *et al*. 2017). The largest *ex situ* collection is held at the International *Musa* Germplasm Transit Centre (ITC) in Leuven, Belgium, with 1,704 accessions. Tracing the origins of triploid cultivated hybrids remains challenging because their ancestry is likely to be complex, involving multiple hybridisation and backcrossing events (Martin *et al*. 2023; Higgins *et al*. 2024).

Fusarium wilt of banana (FWB) has long affected banana production and is caused by different races of the soil-borne fungal pathogen *Fusarium oxysporum* f. sp. *cubense* (Foc) (Pegg *et al*. 2019). The earliest symptoms of FWB are root and rhizome vascular discolouration, but these cannot be detected in the field without destructive sampling. External symptoms appear only after the pathogen has colonised the rhizome. Typical symptoms include chlorosis, wilting, and skirting of older leaves due to vascular blockage (Ploetz 2006). Leaf collapse at the petiole base is then followed by plant death. Other symptoms include malformation of younger leaves, splitting at the base of the pseudostem, and collapse of older leaves while they are still green. In resistant lines of *Musa acuminata* subsp. *malaccensis*, a GFP-tagged Foc race 4 strain showed restricted movement and no sporulation in the rhizome (Chen *et al*. 2019), providing novel insights into how resistance against this pathogen may operate in *Musa* spp.

An epidemic of Foc race 1 in the Americas devastated production of Gros Michel (AAA), the main export banana during the first half of the 20th century (Ploetz 2015). The soil-borne nature of the disease left infested land unsuitable for Gros Michel cultivation (Ploetz 2006; Pegg *et al*. 2019). Cavendish (AAA) cultivars became the dominant export bananas in the 1960s and 1970s because of their resistance to Foc race 1. As banana plantations expanded in Asia, Cavendish genotypes spread worldwide. From the late 1960s to the 1990s, more aggressive forms of FWB emerged in East Asia and Australia, associated with tropical and subtropical race 4 (TR4 and STR4). The distribution of Foc vegetative compatibility groups (VCGs) in Southeast Asia, the centre of origin of *Musa* spp., supports a co-evolution hypothesis in which TR4 was present in native soils before Cavendish (AAA) plantations were established; these plantations were later abandoned following the rapid spread of the disease between 1990 and 1993 (Ploetz and Pegg 1997; Buddenhagen 2009; Mostert *et al*. 2017). Foc TR4 has since spread beyond Asia and Oceania to the Middle East and Africa (Maymon *et al*. 2020; van Westerhoven *et al*. 2023) and to South America, where it has been reported in Colombia, Peru and Venezuela (García-Bastidas *et al*. 2020; Acuña *et al*. 2022; Mejías Herrera *et al*. 2023) and, most recently, in Ecuador in December 2025 (Pachacama *et al*. 2026). Under tropical growing conditions, TR4 appears to be pathogenic to most AAA cultivars, in addition to cultivars susceptible to races 1 and 2 (Ploetz 2015; Mostert *et al*. 2017; Mintoff *et al*. 2021).

Genetic variation in response to FWB has been reported in wild banana germplasm, some hybrids, and mutant lines in several phenotypic studies (Li *et al*. 2015; Zuo *et al*. 2018; Chen *et al*. 2019; Mintoff *et al*. 2021; Thangavelu *et al*. 2025), reflecting the increased attention prompted by the spread of the pathogen and the projected losses associated with the disease over the last decade (Cook *et al*. 2015; Scheerer *et al*. 2018). To date, several resistance or anti-apoptosis genes have been introduced into elite clonal commercial backgrounds through transgenesis or gene editing (Dale *et al*. 2017; Rocha *et al*. 2021; Harding *et al*. 2025). Resistance to Foc appears to be genotype-specific; several wild and cultivated diploid accessions, including Calcutta 4 (AA), Zebrina (AA), Pisang Lilin (AA), Pisang Jari Buaya (AA) and Tuu Gia (AA), are frequently reported as sources of resistance to race 1 and TR4 (Rocha *et al*. 2021). These TR4-resistant diploids have been used as parents in breeding programmes to generate diploid or tetraploid hybrids. However, these hybrids have not met all export market requirements and therefore have not replaced Cavendish (Ploetz 2015; Thangavelu *et al*. 2020). The *Musa* B genome has been associated with abiotic stress tolerance and contains exclusive miRNA families putatively related to stress tolerance (Davey *et al*. 2013); B subgenomes have also been proposed as a rich source of disease-resistance genes (Xie *et al*. 2024). BB germplasm has rarely been screened for response to TR4, and when tested the responses of BB accessions have been inconsistent across studies (Zuo *et al*. 2018; Thangavelu *et al*. 2025). Still, seven plantains (AAB) showed high resistance to TR4 in greenhouse and field experiments (Zuo *et al*. 2018). Additional cultivars carrying a B genome and reported as resistant or tolerant to Foc TR4 include FHIA-21 (AAAB), Plantain number 2 (AAB), Popoulu (AAB), Pisang Nangka (AAA-AAB) and Akpakpak (AAB) (Zuo *et al*. 2018; Thangavelu *et al*. 2025).

Here, we aimed to quantify variation in response to Foc TR4 across a panel of banana accessions selected to represent broad global *Musa* diversity, including major cultivated groups and wild relatives, under controlled-environment conditions. By combining external symptom assessment with image-based quantification of internal corm discolouration, we sought to identify resistant and tolerant germplasm with greater resolution than categorical disease scoring allows. This study’s novelty lies in using a diversity panel structured from previous population structure analysis (Higgins *et al*. 2024) to represent under-evaluated *Musa* groups and in applying quantitative corm phenotyping and mixed-model analysis to compare accessions across experiments. Together, these approaches provide a clearer basis for prioritising resistance sources for breeding and for investigating resistance at the cultivar level rather than inferring it from genome composition alone.

## Materials and methods

### Plant material

Twenty-one banana accessions were used in this study (Table 1): 19 accessions obtained as individual rooted plantlets from the International *Musa* Germplasm Transit Centre (ITC; Bioversity International, Leuven, Belgium) under a Standard Material Transfer Agreement in 2022, and two Cavendish-subgroup lines used as additional TR4 susceptible reference lines, Grande Naine ITC0180 and Tropic Grande Naine, provided by Tropic Biosciences UK Ltd (Norwich, UK). We selected the 19 ITC accessions, as detailed below, based on the population structure of the banana collection at the Colombian genebank (AGROSAVIA) described by Higgins et al. (2024).

**Table 1.** Banana germplasm accessions used in this study, number of plants inoculated and response to Foc TR4. ITC codes, names, genome groups and subgroups follow the ITC passport data supplied with the material. R, resistant; T, tolerant; S, susceptible; VS, very suceptible (see Materials and methods). ^a^The International Musa Germplasm Transit Centre. ^b^Corm data were obtained for 23 plants, of which 22 were analysed after removal of one outlier. ^c^Genome group not recorded in the ITC passport data; assigned to an AAA cluster by Higgins et al. (2024). ^d^Cluster labelled ‘Pome AAAB’ by Higgins et al. (2024). Mock-inoculated plants scored, mock plants with a recorded accession; a further 18 mock-inoculated plants were excluded (see Results).

| **Accession** | **ITC<sup>a</sup> code** | **Biological status** | **Species / genome group** | **Subspecies / subgroup** | **Cluster** (Higgins *et al.* 2024) | **No. of TR4 -inoculated plants scored** | **Mock-inoculated plants scored** | **TR4 response** |
|---|---|---|---|---|---|---|---|---|
| *Musa itinerans* | ITC1526 | Wild | *itinerans* | var. *xishuangbannaensis* | – | 3 | 0 | R |
| *Musa acuminata* subsp. *malaccensis* | ITC0250 | Wild | *acuminata* (AA) | subsp. *malaccensis* | – | 2 | 1 | S |
| *Musa ornata* | ITC0370 | Wild | *ornata* | subsp. *ornata* | – | 9 | 1 | T |
| *Musa schizocarpa* | ITC0926 | Wild | *schizocarpa* (SS) | – | – | 8 | 5 | R |
| *Musa textilis* | ITC1072 | Wild | *textilis* (TT) | subsp. *textilis* | – | 13 | 2 | S |
| Figue Sucrée | ITC0107 | Cultivar | AA | Sucrier | Cl01_Sucrier_AA | 9 | 2 | S |
| Dwarf Cavendish | ITC0002 | Cultivar | AAA | Cavendish | Cl02_Cavendish_AAA | 24<sup>b</sup> | 4 | S (control) |
| Gros Michel | ITC1122 | Cultivar | AAA | Gros Michel | Cl03_Gros_Michel_AAA | 8 | 1 | S |
| IITA hybrid 2145/1320 | ITC1448 | Improved cultivar | Unknown<sup>c</sup> | – | Cl04_unknown_AAA | 8 | 6 | S |
| Green Red | ITC0485 | Cultivar | AAA | Red | Cl05_Red_AAA | 9 | 0 | S |
| FHIA-17 | ITC1264 | Improved cultivar | AAAA | Synthetic tetraploid | Cl06_AAAA | 9 | 2 | S |
| Guineo | ITC0005 | Cultivar | AAA | Mutika/Lujugira | Cl07_Mutika_AAA | 8 | 4 | S |
| Pome | ITC0076 | Cultivar | AAB | Pome | – | 8 | 3 | VS |
| FHIA-02 | ITC0505 | Improved cultivar | AAAB | Synthetic tetraploid | Cl08_Pome_AAAB<sup>d</sup> | 8 | 2 | R |
| FHIA-21 | ITC1332 | Improved cultivar | AAAB | Synthetic tetraploid | Cl09_Africa_AAAB | 6 | 0 | R |
| Mai’a Maoli Koa’e | ITC1167 | Cultivar | AAB | Maia Maoli/Popoulu | Cl10_Popoulu_AAB | 8 | 4 | T |
| Curare | ITC1165 | Cultivar | AAB | Plantain | Cl11_Plantain_AAB | 3 | 0 | R |
| Silver Bluggoe | ITC0364 | Cultivar | ABB | Bluggoe | Cl12_Bluggoe_ABB | 10 | 2 | S |
| Pelipita | ITC0396 | Cultivar | ABB | – | Cl13_Pelipita_ABB | 6 | 1 | T |
| Grande Naine ITC0180 | ITC0180 | Cultivar | AAA | Cavendish | – | 10 | 4 | VS |
| Tropic Grande Naine | – | Cultivar | AAA | Cavendish | – | 11 | 3 | S |

ITC codes, names, genome groups and subgroups follow the ITC passport data supplied with the material. R, resistant; T, tolerant; S, susceptible; VS, very susceptible (see Materials and methods). <sup>a</sup>The International Musa Germplasm Transit Centre. <sup>b</sup>Corm data were obtained for 23 plants, of which 22 were analysed after removal of one outlier. <sup>c</sup>Genome group not recorded in the ITC passport data; assigned to an AAA cluster by Higgins et al. (2024). <sup>d</sup>Cluster labelled ‘Pome AAAB’ by Higgins et al. (2024). Mock-inoculated plants scored, mock plants with a recorded accession; a further 18 mock-inoculated plants were excluded (see Results).

Higgins et al. (2024) identified thirteen clonal clusters in a panel of 188 banana and plantain accessions: nine cultivar groups named according to the passport data stored in the *Musa* Germplasm Information System (MGIS; Ruas *et al*. 2017), one group of unknown AAA accessions lacking passport information, and three groups of synthetic tetraploids from breeding programmes. That study also described wild *Musa* species and thirty-four accessions not assigned to any cluster. From this population structure, we selected one representative from each of the thirteen clusters, plus five wild species, and sourced them from ITC. An accession of the Pome subgroup (AAB) was also included to clarify the differences between the FHIA-02 accession classified as AAAB in Higgins et al. (2024) and the FHIA-02 accessions classified as AAAA in MGIS. When selecting a representative accession for each group, we prioritised accessions duplicated in both the Colombian genebank (AGROSAVIA; Higgins et al. 2024) and the ITC collections; if missing, we selected another member of the same genetic group; or, failing that, an accession of the same subgroup and ploidy. ITC codes, genome groups and subgroups follow the ITC passport data supplied with the material (Table 1).

### Tissue culture and micropropagation

Upon arrival from the ITC, plants were left in the vessels they were shipped in for a recovery period. After acclimatisation, the plants were transferred to rooting medium (RM). Individual rooted plants were later placed in tissue culture vessels containing RM and kept in a growth cabinet at 25°C, 80% relative humidity, and a 16 h light/8 h dark photoperiod until they were ready for subsequent refreshment, micropropagation, or transfer to soil. RM included Murashige and Skoog salts (MS salts), vitamins and a gelling agent. To increase the number of plants, micropropagation was performed using propagation medium (PM; Fig. 1A–E). Plants were cleaned of dead tissue and trimmed, and explants were divided and transferred to culture jars containing PM. New propagation cycles can be started as soon as two weeks or as late as three months after micropropagation, depending on the plant’s growth rate and experimental needs.

**Figure 1.**
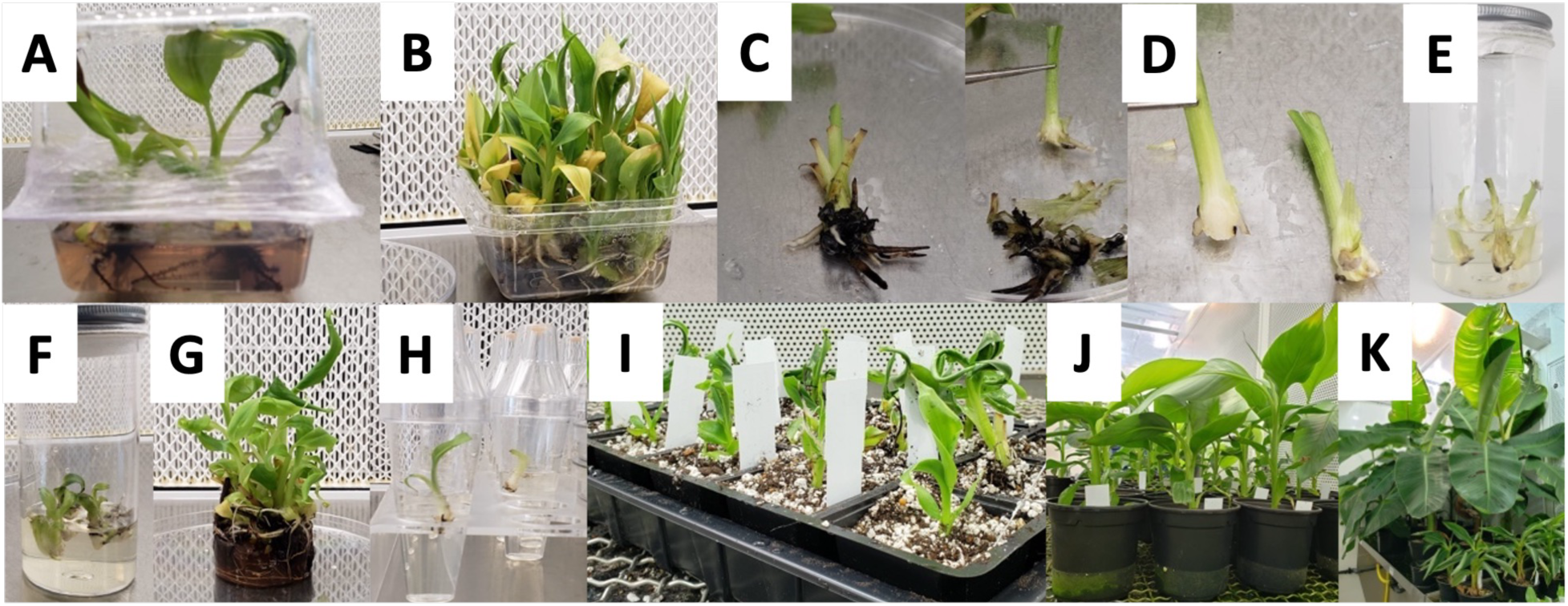
Banana plants were micropropagated and established in soil before inoculation. (A–E) Tissue culture and micropropagation: (A) plantlets as received from ITC; (B) plantlets transferred to a dish; (C) individual plants separated and cleaned of dead tissue; (D) plants prepared for division; (E) explants placed in a jar of propagation medium (PM) to start a micropropagation cycle. (F–K) Plant refreshment and soil establishment: (F, G) plants removed from PM jars and cleaned of dead tissue and medium; (H) individual plants cultured in rooting medium (RM) until transfer to the growth cabinet; (I) plants transferred to trays with soil mix and covered during the initial acclimatisation period to maintain high humidity; (J) plants in pots, ready for inoculation assays; (K) plants re-potted when larger plants are required. Alt text: Eleven photographs showing the steps of banana tissue culture, micropropagation and transfer of plantlets to soil trays and pots.

### Soil establishment

Plants were refreshed in vitro into new RM at least one week before transfer from RM to soil to allow for root recovery. Only plants that had at least two roots longer than 1 cm and three leaves were transferred. Plants were transferred to a sterile soil mix (JIC barley mix: 60% peat, 20% grit/sand, and 20% perlite) in cell trays, then kept in a controlled-environment cabinet under controlled temperature, photoperiod and high relative humidity (Fig. 1F–K). Plants were irrigated by automatic flooding and were covered during the initial acclimatisation period. After acclimatisation, the plants were transplanted into pots with the same soil mix and grown for three weeks before TR4 inoculation assays.

The composition of the tissue-culture media (RM and PM), together with specific details of the associated micropropagation, culture and acclimatisation procedures, is proprietary to Tropic Biosciences UK Ltd (Norwich, UK) and is therefore not disclosed. All other materials and methods are described in full.

### Foc TR4 inoculum

Inoculum was produced from Foc TR4 isolate UK0001, recovered from a symptomatic plant at the Eden Project (Cornwall, UK) and sequenced by Warmington et al. (2019). Genomic analyses place this isolate close to the Indonesian TR4 isolate II-5 at the sequence level (Warmington *et al*. 2019). For inoculum production, 20 µL of a −80°C stock of isolate UK0001 was plated onto potato dextrose agar (PDA) and incubated for 7 days at 25°C in the dark. A 5 mm plug was transferred onto a new PDA plate and incubated for a further 7 days. Three plugs from this second plate were added to 120 mL of potato dextrose broth (PDB) and incubated in a water-bath shaker at 25°C and 180 rpm for 5 days. The liquid culture was filtered through sterile cheesecloth to remove mycelial fragments. The filtrate was divided evenly between three 50 mL Falcon tubes and centrifuged for 10 minutes at 3,000 rpm at 20°C. The pellet in each tube was rinsed twice in 40 mL of distilled water and centrifuged for 5 minutes at 3,000 rpm. After the second wash, the pellet in the first tube was resuspended in 15 mL of distilled water and transferred to the next 50 mL tube, repeating the resuspension until all inoculum was combined. A 0.5 mL aliquot from each tube was taken for quality control, along with the mock PDB control. Finally, all the TR4 inoculum was pooled, and a 0.5 mL aliquot was taken for spore counting, using a Neubauer chamber. Inoculum was prepared at 2 × 100 spores/mL, with 200 mL applied per plant.

The control inoculum for the mock-inoculations was prepared in the same way, using TR4-free PDB incubated alongside the inoculum preparations.

### Experimental design and inoculation

Inoculation assays were conducted in a controlled-environment cabinet as six independent experiments between 2023 and 2025, numbered 1 and 3–7; a further experiment (experiment 2) was lost to a cabinet failure and no data from it were used. Each experiment was treated as a block, and the four internal quadrants of the cabinet (A–D) were treated as replicates. Because the number of plants available per accession was limited by micropropagation, each experiment included a subset of the accessions, giving an incomplete block design in which each accession other than Dwarf Cavendish was tested in two to four experiments. Within each experiment, TR4-inoculated plants of each accession were distributed across the quadrants alongside mock-inoculated plants of the same accessions. Cultivar Dwarf Cavendish was included as the susceptible control in every experiment and quadrant, to allow responses to be compared among experiments and quadrants. In total, 180 plants were inoculated with Foc TR4 and 65 plants were mock-inoculated; the 47 mock-inoculated plants with a recorded accession are listed in Table 1. All accessions were tested in a minimum of two independent inoculation experiments.

Five days before inoculation, plants were moved onto individual trays and placed in the cabinet, and water was withheld before inoculation. On the day of inoculation, the number of open leaves and, in some experiments, plant height were recorded (Table S1). Control plants were mock-inoculated first, followed by TR4-inoculated plants, to minimise the risk of cross-contamination. Inoculation involved gently pouring 200 mL of mock solution or spore suspension (2 × 10^5^ spores/mL) around the base of the pseudostem to maximise contact with the roots (García-Bastidas *et al*. 2019). From one week after inoculation, plants were watered from below with 200 mL of water per tray twice weekly, until the end of the assay or until plant death.

### External and internal scoring

We scored external symptoms weekly from inoculation (week 0) to six weeks after inoculation (wpi) by counting the total number of open leaves and the number of yellow (wilting) leaves. Leaves were recorded as yellow when more than 50% of their area was discoloured. Plant height was also recorded at inoculation (week 0) and at 6 wpi, and pseudostem splitting was noted, in a subset of experiments (Table S1). At 6 wpi, each plant was photographed before internal symptoms were assessed. For this, the pseudostem was cut 10 cm above the corm, the roots were removed and the soil was washed away with water; the remaining stem was then cut longitudinally to expose the inner corm tissue, and both halves were photographed with a 5 cm label for scale and colour reference. Camera, lighting, and focal distance were standardised across experiments. Using ImageJ (Schneider *et al*. 2012), the total corm area was manually outlined and measured, and the healthy (non-discoloured) area was then delimited using the colour contrast between healthy and discoloured tissue and measured (Table S2). The percentage of healthy corm area (100 × healthy area/total corm area) was calculated for each of the two halves of each corm, and completely discoloured corms were recorded as 0% healthy area. Corms of mock-inoculated plants were sectioned and photographed in the same way.

### Data analysis

All analyses were performed in R (R Core Team 2024), and figures were drawn with ggplot2 (Wickham 2016). For external symptoms, TR4-inoculated plants were grouped by the response class of their accession (R, T, S or VS; see below), and the total number of leaves and the percentage of yellow leaves were each analysed with a single linear mixed model fitted to all weekly observations (weeks 0–6), with group (the four TR4 groups and mock-inoculated plants), week and their interaction as fixed effects and block (experiment × quadrant), accession and plant as random effects. Plants that died during experiments were assigned their last recorded number of leaves and 100% yellow leaves for the remaining weeks. Within each week, each TR4 group was compared with the mock-inoculated plants using Dunnett’s test (Table S3). Height gain from inoculation to 6 wpi was analysed with a linear mixed model including group as a fixed effect and block and accession as random effects. The proportion of plants with pseudostem splitting is shown with Wilson 95% confidence intervals (Fig. S1).

For the percentage of healthy corm area, data from TR4-inoculated plants were analysed as an incomplete block design using best linear unbiased estimates (BLUEs) from a linear mixed model fitted with lme4 (Bates *et al*. 2015) and lmerTest (Kuznetsova *et al*. 2017), in which accessions (treatments) were fitted as fixed effects and replicates (quadrants) and blocks (experiments) within replicates as random effects: *y_ijk_ = m + t_i_ + :r_j_ + b_jk_ + e_ijk_*, where, *y_ijk_* is the percentage of healthy corm area measured on a corm half of the *i^th^* accession in the *j^th^* replicate (quadrant) within the *k^th^* block (experiment), *m* is the overall mean, *t_i_* is the fixed effect of the accession, *:r_j_* is the random effect of the replicate (cabinet quadrant), *b_jk_* is the random effect of the block (experiment) within the replicate, and *e_ijk_* is the residual error. The null hypothesis was that all accession effects were equal *(H_O_: t_1_ = t_2_ = - = t_p_)*, while the alternative hypothesis was that at least one accession effect differed. The model was fitted in R as pct_healthy ∼ accession + (1 | quadrant) + (1 | quadrant:experiment). Model assumptions were checked using residual plots for normality and homogeneity of variance. BLUEs and contrasts were obtained with emmeans (Lenth 2024), using the Kenward–Roger approximation for the degrees of freedom (Halekoh and Højsgaard 2014) (Table S4). Each accession was compared with Dwarf Cavendish, the susceptible control included in every experiment, using Dunnett’s test (Table S5), and all pairs of accessions were compared using Tukey’s test (Table S6).

Accessions were assigned to response classes from the corm data. Accessions whose BLUE was significantly higher than that of Dwarf Cavendish (Dunnett’s test, *P* < 0.05) were classified as resistant (R) when all their corm halves had more than 80% healthy area, and as tolerant (T) otherwise. Accessions whose BLUE was significantly lower than that of Dwarf Cavendish were classified as very susceptible (VS), and all remaining accessions as susceptible (S).

## Results

Twenty-one accessions (Table 1) were compared across six TR4 inoculation experiments conducted from 2023 to 2025. Poor propagation resulted in fewer plants being tested for four accessions: *M. acuminata* subsp. *malaccensis* (two plants in two experiments), *M. itinerans* (three plants in two experiments), Curare (Plantain subgroup, AAB; three plants in two experiments) and Pelipita (ABB; six plants in three experiments). Missing replicates in blocks were accounted for in the mixed model used for data analysis. Of the 65 mock-inoculated plants, 47 were included in the analysis of external symptoms; the remaining 18 were excluded because the plant died for reasons unrelated to the inoculation or their accession was not recorded.

### Internal scoring of TR4 symptoms

Longitudinal sections of the corm at 6 wpi revealed contrasting internal symptoms among accessions, ranging from no discolouration in accessions such as *M. schizocarpa* to extensive discolouration in accessions such as Figue Sucrée (Fig. 2A). The percentage of healthy corm area was quantified for both halves of each corm. Mock-inoculated corms showed no discolouration (Fig. S2). All corm halves of FHIA-02, FHIA-21, *M. itinerans*, *M. schizocarpa* and Curare had more than 80% healthy area, whereas more than half of the corm halves of Pome, Figue Sucrée, Grande Naine ITC0180 and FHIA-17 had 5% or less healthy area (Fig. S3).

**Figure 2.**
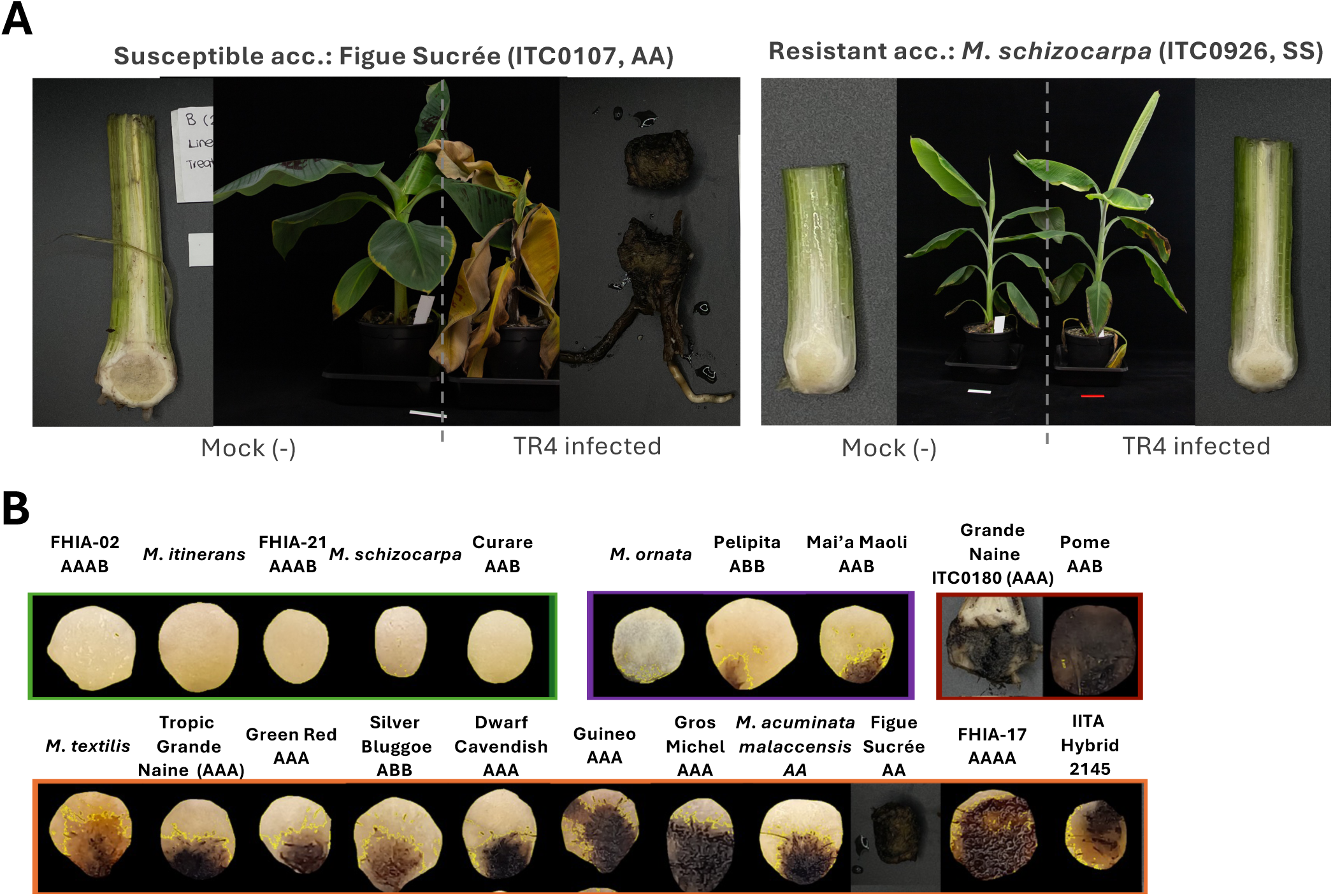
Corm discolouration after Foc TR4 inoculation varies widely among accessions. Internal symptoms of Foc TR4 infection at 6 weeks after inoculation. (A) Mock- and TR4-inoculated plants and longitudinal corm sections of a susceptible accession (Figue Sucrée, ITC0107, AA) and a resistant accession (*M. schizocarpa*, ITC0926, SS). (B) Representative corm halves of a TR4-inoculated plant of each accession, framed by response class: resistant (green), tolerant (purple) and very susceptible (red) in the upper row, and susceptible (orange) in the lower row. Yellow outlines show the healthy area delimited in ImageJ. Alt text: Photographs of banana plants and corm sections showing healthy corms in a resistant wild species and dark, discoloured corms in a susceptible cultivar, followed by one corm section for each of the 21 accessions.

We estimated BLUEs of the percentage of healthy corm area with experiments (blocks) and cabinet quadrants (replicates) treated as random effects, to account for variation due to the experimental design (Table S4). Healthy corm area in the susceptible control varied among experiments, with a mean percentage of healthy corm area per experiment ranging from 15% to 59% for Dwarf Cavendish (Fig. 3A; Fig. S4), supporting the use of a data analysis model that accounts for experiment effects. The residuals of the mixed model were approximately normally distributed (Fig. S5A) and evenly distributed above and below zero across fitted values (Fig. S5B); the deviations in the tails and the diamond-shaped pattern near 0% and 100% reflect the bounded nature of the response variable.

**Figure 3.**
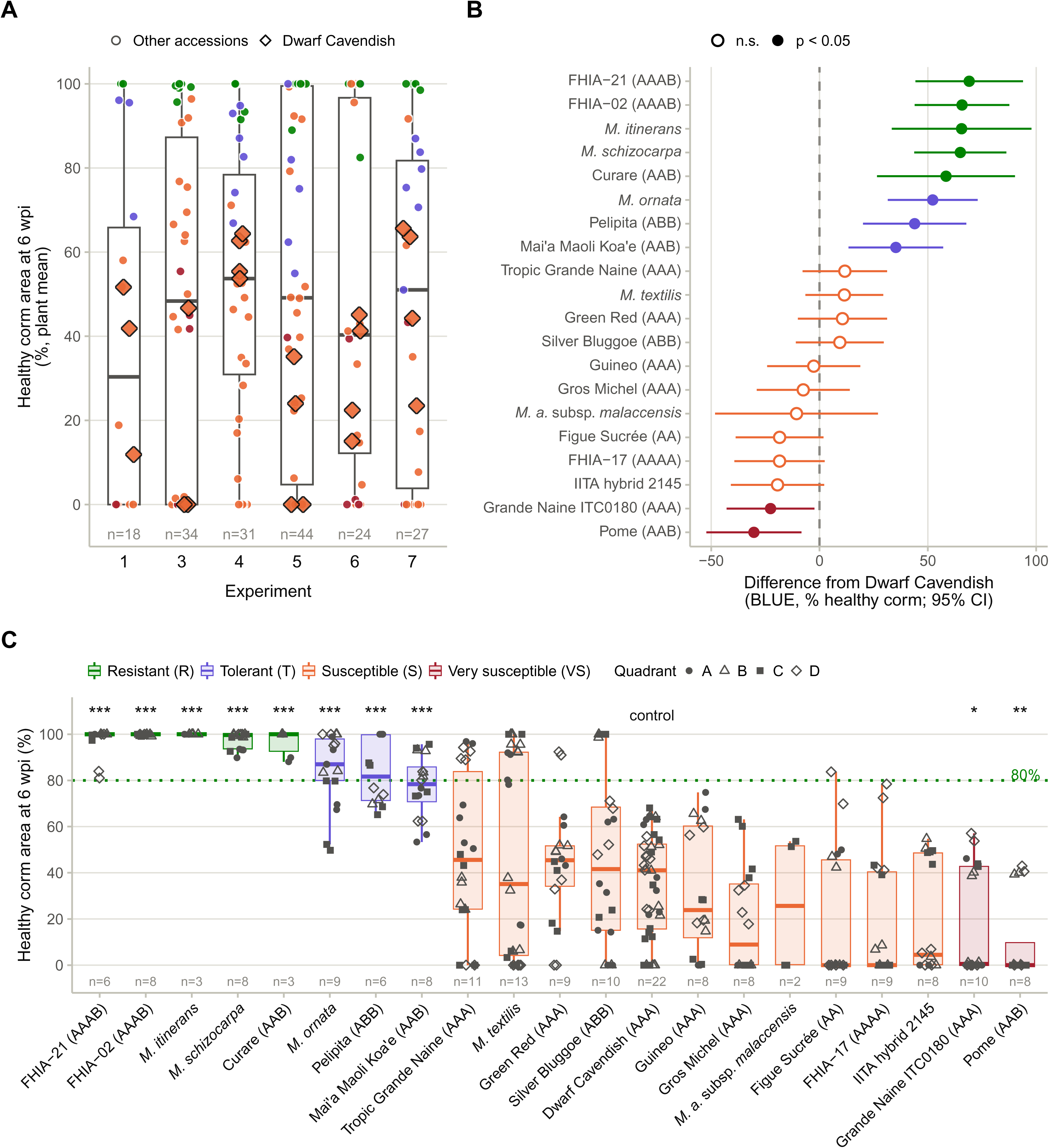
Eight accessions keep significantly more healthy corm tissue than Dwarf Cavendish. Internal symptoms of Foc TR4 infection quantified as the percentage of healthy corm area at 6 weeks after inoculation (wpi). (A) Percentage of healthy corm area (plant means) of all TR4-inoculated plants in each experiment; diamonds, Dwarf Cavendish; circles, other accessions, coloured by response class; n, number of plants. (B) Difference between the best linear unbiased estimate (BLUE) of each accession and that of the susceptible control Dwarf Cavendish, with 95% confidence intervals; filled symbols, significant differences (Dunnett’s test, *P* < 0.05); open symbols, not significant. (C) Percentage of healthy corm area of each corm half by accession; symbols indicate the cabinet quadrant (A–D), boxes are coloured by response class (R, resistant; T, tolerant; S, susceptible; VS, very susceptible) and n is the number of plants. Asterisks indicate significant differences from Dwarf Cavendish (Dunnett’s test; \**P* < 0.05, \*\**P* < 0.01, \*\*\**P* < 0.001). The green dotted line marks 80% healthy corm area, the threshold used to classify resistant accessions. Boxes (A, C) show the median and interquartile range, and whiskers extend to 1.5 times the interquartile range. Alt text: Forest plot and box plots of the percentage of healthy corm area by accession and by experiment; eight accessions have much higher healthy corm area than the Dwarf Cavendish control, and two have lower.

The differences between the BLUE of each accession and that of Dwarf Cavendish, the control, with 95% confidence intervals, are shown in Fig. 3B. Accessions with positive values were more resistant to TR4 than the control, and those with negative values were more susceptible. The BLUE of Dwarf Cavendish was 34.8% healthy corm area. Eight accessions had significantly higher percentages of healthy corm area than Dwarf Cavendish (Dunnett’s test, *P* < 0.001; Fig. 3B; Table S5), with differences of 35 to 69 percentage points, and Tukey pairwise comparisons did not show significant differences among these eight accessions (Table S6). Five of them, FHIA-02, FHIA-21, *M. itinerans*, *M. schizocarpa* and Curare, were classified as resistant (R), because all their corm halves had more than 80% healthy area (Fig. 3C). The other three, *M. ornata*, Pelipita and Mai’a Maoli Koa’e, were classified as tolerant (T), because some of their plants had less than 80% healthy corm area (minimum plant means of 51–67%). Two accessions, Grande Naine ITC0180 (AAA) and Pome (AAB), had significantly lower percentages of healthy corm area than Dwarf Cavendish (*P* < 0.05 and *P* < 0.01, respectively; Fig. 3B, C) and were classified as very susceptible (VS); 60% and 75% of their corm halves, respectively, had 5% or less healthy area (Fig. S3). The remaining 11 accessions did not differ significantly from Dwarf Cavendish and were classified as susceptible (S).

The response class of each accession is summarised in Table 1. The resistant and tolerant accessions comprised three wild species with neither A nor B genomes (*M. itinerans*, *M. schizocarpa* and *M. ornata*) and five accessions with at least one B genome (FHIA-02 and FHIA-21, AAAB; Curare and Mai’a Maoli Koa’e, AAB; Pelipita, ABB). All accessions with only A genomes (AA, AAA and AAAA), and the IITA hybrid 2145 assigned to an AAA cluster, were susceptible or very susceptible.

### External scoring of TR4 symptoms

External symptoms were scored weekly for six weeks as the number of open leaves and the percentage of yellow leaves (Fig. 4). For this analysis, TR4-inoculated plants were grouped by the response class of their accession, determined from the internal symptoms (see above): resistant (R; 28 plants of five accessions), tolerant (T; 23 plants of three accessions), susceptible (S; 111 plants of 11 accessions) and very susceptible (VS; 18 plants of two accessions). Resistant and tolerant accessions had significantly less corm discolouration than the susceptible control Dwarf Cavendish, and very susceptible accessions significantly more (see Internal scoring of TR4 symptoms). These were compared with leaf development and health of mock-inoculated plants. The total number of leaves increased steadily over the six weeks and was similar in all TR4-response groups and in mock plants, except plants of VS accessions which had significantly more leaves following TR4-inoculation than mock-inoculated plants at week 2 (*P* = 0.03; Fig. 4A; Table S3). In contrast, plants of S and VS accessions had a significantly higher percentage of yellow leaves than mock plants from week 4 onwards (*P* < 0.001; Fig. 4B), reaching on average about 55% and 80% yellow leaves, respectively, at week 6, compared with about 30% in mock plants. Plants of R and T accessions did not differ significantly from mock plants at any week (Fig. 4B). Leaf yellowing and wilting are typically observed in banana plants following TR4 infection because the fungus colonises the xylem and disrupts the normal growth of the plant. Dwarf Cavendish (AAA) showed an atypical progression of symptoms following TR4-inoculation, with the number of new leaves comparable to or higher than in mock-inoculated plants. However, these new leaves emerged more quickly and were stunted and deformed, with a small surface area. New leaves emerged in rapid succession from the same position on the pseudostem, which did not elongate between successive leaves, so the leaves remained clustered at the top of a short pseudostem. In plants of resistant accessions, the rate of leaf emergence was similar between TR4- and mock-inoculated plants, and yellow leaves, when present, were attributable to senescence, as they were observed at a similar rate in both treatments.

**Figure 4.**
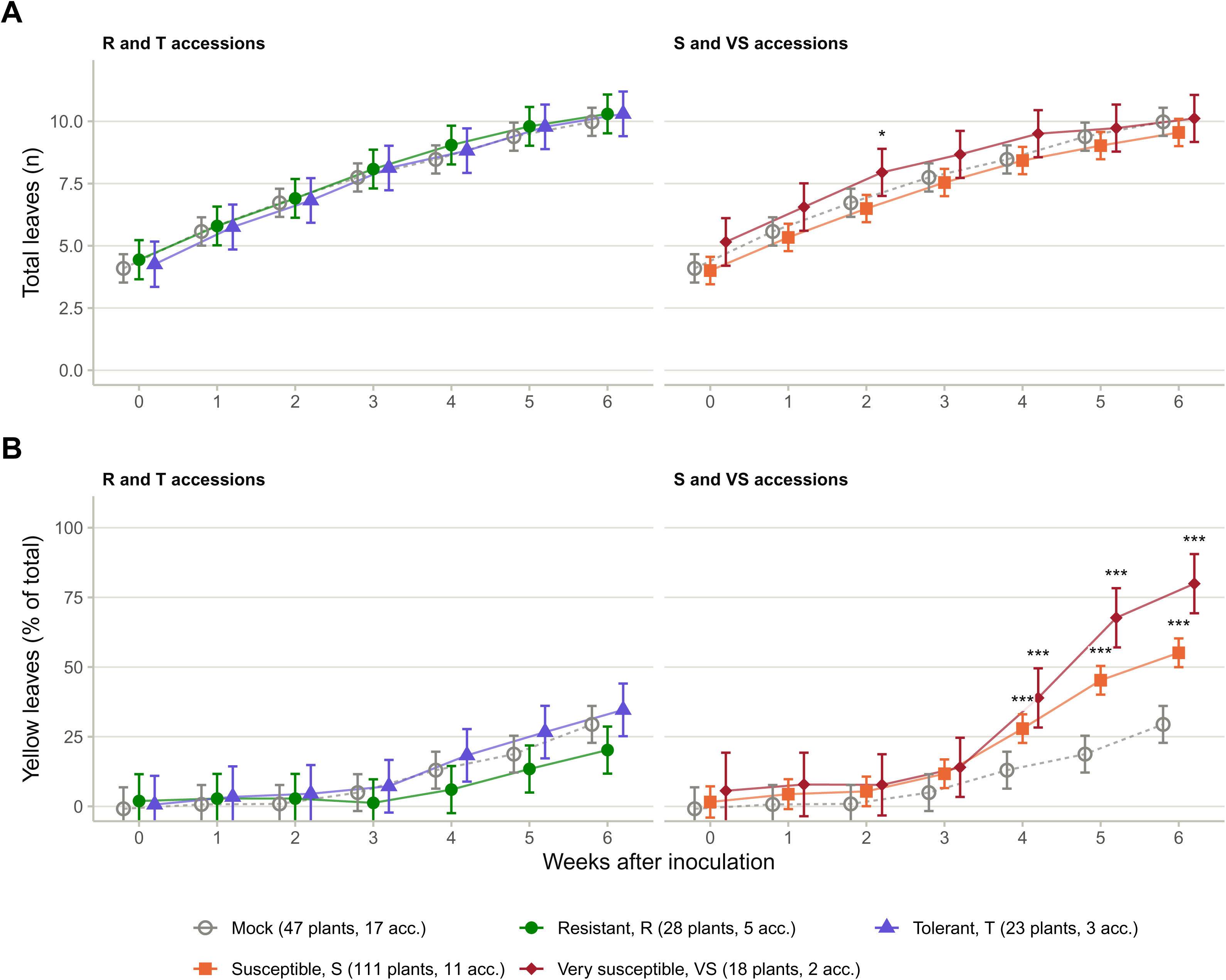
Leaf yellowing rises from 4 wpi in susceptible accessions but not in resistant or tolerant ones. External symptoms after inoculation with Foc TR4. Adjusted means of (A) the total number of leaves and (B) the percentage of yellow leaves from inoculation (week 0) to 6 weeks after inoculation, for TR4-inoculated plants grouped by the response class of their accession (left, resistant (R) and tolerant (T) accessions; right, susceptible (S) and very susceptible (VS) accessions) and for mock-inoculated plants (grey open circles and dotted line, shown in both panels). Means were estimated with a single linear mixed model for each trait fitted to all weekly observations, with group, week and their interaction as fixed effects and block (experiment × quadrant), accession and plant as random effects (Table S3); error bars, 95% confidence intervals. Asterisks indicate significant differences from mock plants within a week (Dunnett’s test; \**P* < 0.05, \*\*\**P* < 0.001). The numbers of plants and accessions in each group are given in the key. Data from experiments 1 and 3–7. Alt text: Line plots of total leaves and percentage of yellow leaves over six weeks after inoculation; susceptible and very susceptible groups have a higher percentage of yellow leaves than mock plants from week 4, whereas resistant and tolerant groups resemble mock plants.

Height gain from inoculation to 6 wpi did not differ significantly between any TR4-response group and mock-inoculated plants (*P* ≥ 0.24), although plants of S and VS accessions gained on average about 13 and 11 cm, compared with about 16 cm in mock plants (Fig. S1A). Pseudostem splitting was rare (4.3% of evaluated TR4-inoculated plants) and was not observed in mock-inoculated plants or in plants of R accessions (Fig. S1B).

## Discussion

This study set out to examine the response to Foc TR4 across a genetically diverse panel of Musa spp., and identified new sources of resistance within it. The accessions assessed represented, to a large extent, the primary and tertiary diversification of *Musa* spp., including some of the most common and widely grown cultivars worldwide: nine cultivar subgroups recorded in MGIS plus Pelipita, three synthetic tetraploids and an IITA hybrid, together with five wild species. We obtained the accessions representing each cluster from ITC, whereas the full diversity panel had previously been sequenced from the AGROSAVIA collection in Colombia (Higgins *et al*. 2024).

Four of the five major genomic groups were represented: AA, AAA, AAB and ABB, with only the AB hybrids absent. No wild *M. balbisiana* (BB) accession was included. AB interspecific diploids are under-represented in germplasm collections because they arose through secondary diversification in regions where the natural distributions of the wild progenitor species overlap, mainly in South Asia (Fuller and Madella 2009). Nevertheless, the absence of interspecific diploids does not imply that their genetic background is absent from the panel. Analysis of AB homologous exchanges supported contributions from AB diploids to the Bluggoe (ABB) and Pelipita (ABB) clusters, both of which were included (Cenci *et al*. 2021). Including *M. schizocarpa* (S genome donor) was also important, as this species contributed to many cultivar groups, though to varying extents, as supported by chromosome painting and Relative Averaged Alignment studies (Martin *et al*. 2023; Higgins *et al*. 2024). The presence of S ancestry across a range of genome compositions provides evidence of early recombination between *M. acuminata* and *M. schizocarpa* before the establishment of cultivated AAA, AAB, and ABB cultivars (Heslop-Harrison and Schwarzacher 2007; Li *et al*. 2013).

All accessions with only A genomes (AA, AAA and AAAA) were susceptible or very susceptible to TR4, including the well-known susceptible cultivars Dwarf Cavendish (AAA) and Gros Michel (AAA). In this study, the *M. acuminata* subsp. *malaccensis* (AA) accession tested appeared susceptible (two plants), although other genotypes of this wild sexual species have been reported as resistant to Foc TR4 (Dale *et al*. 2017; Chen *et al*. 2019). Natural variation in TR4 responses is expected in sexually reproducing taxa where recombination occurs. Resistance to TR4 has already been experimentally validated within the A genome: the *RGA2* gene, isolated from a resistant *M. acuminata* subsp. *malaccensis* genotype, conferred resistance when introduced into a Cavendish background, and the resulting transgenic line has recently been approved for commercial production and consumption in Australia (Dale *et al*. 2017; Harding *et al*. 2025). Similarly, the *M. textilis* accession tested here was susceptible, although its response was bimodal, with five of 13 plants having more than 80% healthy corm area and three having none. Natural variation in response to TR4 might therefore be expected in this species, which originates from the Philippines, where TR4 is present (Mostert *et al*. 2017); to our knowledge, no resistant genotype of this species has yet been reported. Other susceptible accessions included Silver Bluggoe (ABB), IITA hybrid 2145 and Pome (AAB). FHIA-02 (AAAB) represented a cluster labelled ‘Pome AAAB’ by Higgins et al. (2024), because one of its members was recorded as Pome. Notably, FHIA-02 was resistant, whereas the Pome (AAB) accession tested here was among the most susceptible accessions in our study (Fig. 3).

The presented results suggest that the B genome may represent an additional source of resistance to TR4. Five of the eight resistant or tolerant accessions contained at least one B genome, and five of the seven B-containing accessions tested were resistant or tolerant, compared with none of the accessions with only A genomes. The association is therefore one of frequency rather than sufficiency: Pome (AAB) and Silver Bluggoe (ABB) carried a B genome yet were among the most susceptible accessions in the panel, and three of the eight resistant or tolerant accessions carried neither an A nor a B genome. This interpretation has limitations. No *M. balbisiana* (BB) or AB accession was tested. Three of the eight resistant or tolerant accessions were wild species with neither A nor B genomes, and the two resistant FHIA tetraploids derive from breeding programmes that used improved diploid parents selected for disease resistance, so part of their resistance may derive from A-genome donors. Moreover, *M. balbisiana* (BB) and other wild diploids have not been studied as extensively as A-genome accessions. One wild *M. balbisiana* accession was highly susceptible to TR4, in contrast to five resistant wild *Musa* spp., after inoculation with a locally sourced TR4 strain in China (Li *et al*. 2015). Two independent studies tested the accession Tani BB (ITC1120) against TR4 under glasshouse and field conditions, using the same inoculation and scoring methodology. Zuo et al. (2018) initially reported Tani BB as resistant in the glasshouse and immune in the field. However, the same accession was later reported by Thangavelu et al. (2025) as moderately resistant in both glasshouse and field conditions. Given these differing outcomes, we noted that the Foc TR4 isolate and inoculum concentration used in the glasshouse inoculations differed between the two studies. Moreover, the field experiments did not report the Foc TR4 isolate or concentration, as the study used soils previously contaminated with TR4. It is also unclear how disease scores were translated into resistance categories, and the raw scoring data were not made available (Thangavelu *et al*. 2025). Differences in response to TR4 reported even for the same clone across studies highlight the challenge of reproducibility in inoculation assays. Zuo et al. (2018) reported Pisang Ceylan AAB (ITC1441) as resistant, with no infection detected in the rhizome of field-grown plants after two years and Foc TR4 confirmed by PCR in infected tissues. However, the same accession (ITC1441) was later reported by Zhan et al. (2022) as showing intermediate resistance. Both studies evaluated the same clone and the same Foc TR4 strain in the glasshouse experiments, and the plants were grown in the same field in China. Furthermore, the other three accessions shared between the two studies showed similar responses to Foc TR4 (Zuo *et al*. 2018; Zhan *et al*. 2022).

We observed variation in the TR4 response of accessions across inoculation experiments under our bioassay conditions. The effects of the experimental design were accounted for by estimating BLUEs from a model that treated accession effects as fixed while accounting for the random effects of the incomplete experimental blocks and of the quadrants within the cabinet. This was important because the response of the susceptible control varied considerably among experiments (Fig. 3A). Including Dwarf Cavendish in every experiment (22 plants analysed in total) provided a common reference against which all accessions could be contrasted, even though each of the other accessions was tested in only two to four experiments. The assays used young plants under controlled conditions, with an endpoint of 6 wpi, so slower-developing infections may have been under-scored, and responses in the field may differ. In addition, we classified *M. itinerans* and Curare as resistant based on three plants each, although the same *M. itinerans* accession (ITC1526) has also been rated highly resistant elsewhere (Zuo *et al*. 2018; Rouard *et al*. 2026).

Several of our results agree with previous screenings of the same or related accessions. The *M. itinerans* accession tested here (ITC1526) has been rated highly resistant to TR4 (Zuo *et al*. 2018), and FHIA-02 has been reported as resistant to TR4 in glasshouse and field studies (Chen *et al*. 2019; Mintoff *et al*. 2021); both ITC accessions are included in a recently curated subset of TR4-resistant germplasm (Rouard *et al*. 2026). Most plantains show high resistance to TR4 (Zuo *et al*. 2018; Zhan *et al*. 2022), consistent with Curare’s resistance. In contrast, accessions of the same subgroup have sometimes given contrasting results: two accessions of the Maia Maoli/Popoulu subgroup differed in their response to TR4 (Zhan *et al*. 2022), Pelipita accessions have been rated from highly resistant to susceptible (Rouard *et al*. 2026), and Pome-subgroup accessions were resistant in field and glasshouse trials in China (Zhan *et al*. 2022), whereas the Pome accession tested here (ITC0076) was very susceptible. Such variation within subgroups, together with the variable responses of Mai’a Maoli Koa’e and Pelipita in our study, supports investigating resistance at the cultivar level.

Further studies of AAB/ABB cultivars and AB hybrids will be needed to clarify the role of the B genome in TR4 resistance. The source of resistance should be investigated at the cultivar level rather than at the genomic-group level, as variation within groups has already been observed. Variation in response to TR4 at the cultivar level is likely associated with specific combinations of genes and regulatory elements involved in resistance, such as NLR immune receptors, of which RGA2 is one example (Dale et al. 2017). Characterisation of NLRs across the *Musa* genus should therefore be a priority for future research. There is also a need for a robust and reproducible scoring system for TR4 bioassays that is less prone to subjective assessment and to the resulting loss of statistical power. In this regard, a continuous measure of corm discolouration, such as the image-based percentage of healthy corm area used here, is preferable to categorical scales. Screening that scores infection on a discrete scale rather than as a quantitative trait reduces a continuous response to broad categories, which discards information and reduces statistical power. Categorical scales may also saturate: the corms of Dwarf Cavendish plants in our study ranged from 0% to 66% healthy area, a range that would fall within the two most severe classes of a typical six-class rhizome discolouration scale. Categorical scoring also makes comparison across studies more difficult, particularly when the raw scores are not made available.

## Conclusions

The quantitative phenotyping used here revealed substantial variation in response to Foc TR4 across a panel designed to capture understudied *Musa* diversity. Five accessions were classified as resistant and three as tolerant, whereas all accessions carrying only A genomes were susceptible or very susceptible in this panel, including well-established susceptible controls. Among them is the M. acuminata subsp. malaccensis accession tested, which was represented by only two plants and for which resistant genotypes have been reported elsewhere; the result therefore reflects the specific accessions assessed rather than the A genome as a whole. Although resistance cannot be assigned to a single genomic group, the frequency of resistant or tolerant responses among accessions carrying at least one B genome suggests that this genomic background may represent an underexplored source of resistance against TR4; two B-containing accessions were nonetheless highly susceptible, so a B genome is not by itself sufficient for resistance. In addition, continuous corm discolouration measurements allowed accessions to be compared across experiments on a common quantitative scale and avoided the limitations associated with categorical disease classes. These findings provide a basis for prioritising germplasm in resistance breeding and for future studies aimed at identifying the genes and regulatory mechanisms underlying TR4 resistance.

## Data Availability

The data supporting the findings of this study, including the weekly external symptom scores per plant, corm measurements and model estimates, are provided as Supporting Information (Tables S1–S6). The R scripts used for the analyses and figures will be deposited in a GitHub public repository (https://github.com/DeVegaGroup/TR4_response_screening).

## Acknowledgements

We thank Leo Wilkinson for assistance with the first inoculation experiment, and Finlay Taylor, Sophie Thomas, and the rest of the pathology and media teams at Tropic Biosciences UK Ltd. We also acknowledge the International Musa Germplasm Transit Centre (Bioversity International, Leuven, Belgium) for providing the germplasm. The authors used Generative AI (Claude, Anthropic) to help check the curated data, reproduce the statistical analyses, and edit the manuscript text. The authors reviewed and verified all outputs and take full responsibility for the content.

## Supporting Information

**Figure S1. Foc TR4 did not significantly reduce height gain, and pseudostem splitting was rare.** Plant height gain and pseudostem splitting after inoculation with Foc TR4. (A) Adjusted mean height gain from inoculation (week 0) to 6 weeks after inoculation, with 95% confidence intervals, for mock-inoculated plants and TR4-inoculated plants grouped by the response class of their accession (R, resistant; T, tolerant; S, susceptible; VS, very susceptible), from a linear mixed model with block and accession as random effects; n, number of plants. (B) Percentage of plants with pseudostem splitting noted, with Wilson 95% confidence intervals; numbers are plants with splitting/plants scored.

**Figure S2. Corm discolouration occurs only in TR4-inoculated plants.** Photographic summary of external and internal symptoms at 6 weeks after inoculation. (A) Representative TR4- and mock-inoculated plants and longitudinal corm sections of a resistant and a susceptible accession; scale bars, 5 cm. (B–D) Corm halves of one TR4-inoculated plant (upper two rows) and one mock-inoculated plant (lower two rows) per accession (one accession per column) for (B) resistant accessions (green frame), (C) tolerant accessions (purple frame) and (D) susceptible and very susceptible accessions (red frame). Yellow outlines show the healthy corm areas delimited in ImageJ.

**Figure S3. All corm halves of resistant accessions stay above 80% healthy area.** Distribution of classes of healthy corm area scored for each accession tested. Bars show, for each accession, the proportion of corm halves of TR4-inoculated plants in each class of healthy corm area at 6 weeks after inoculation; n, number of plants. Accessions are ordered as in Fig. 3C, and the strip above the bars shows the response class (R, resistant; T, tolerant; S, susceptible; VS, very susceptible).

**Figure S4. Healthy corm area of the Dwarf Cavendish control varies between experiments.** Variability of the Dwarf Cavendish control. (A) Percentage of healthy corm area of the susceptible control Dwarf Cavendish in each experiment after removal of one outlier; symbols indicate the cabinet quadrant (A–D). (B) Difference between the BLUE of each accession and that of Dwarf Cavendish, with 95% confidence intervals, coloured by response class; filled symbols, *P* < 0.05 (Dunnett’s test).

**Figure S5. Residuals of the corm mixed model are approximately normal.** Mixed-model diagnostics of TR4 inoculation scoring data. (A) Normal quantile–quantile plot of the residuals of the mixed model for the percentage of healthy corm area. (B) Residuals against fitted values; the diamond-shaped pattern near 0% and 100% reflects the bounded response variable.

**Table S1.** Weekly external symptom scores per plant (total and yellow leaves, plant height and pseudostem splitting).

**Table S2.** Corm area measurements for each corm half at 6 weeks after inoculation.

**Table S3.** External symptoms: adjusted means per week for each TR4-response group and mock-inoculated plants, and Dunnett’s comparisons with mock-inoculated plants.

**Table S4.** Best linear unbiased estimates (BLUEs) of the percentage of healthy corm area for each accession.

**Table S5.** Comparison of each accession with the susceptible control Dwarf Cavendish (Dunnett’s test).

**Table S6.** Pairwise comparisons between accessions (Tukey’s test).

