## Supplementary figure S1 for "Evaluating novel sources of resistance to *Fusarium oxysporum* f. sp. *Cubense* tropical race 4 (TR4) in a global banana diversity panel"

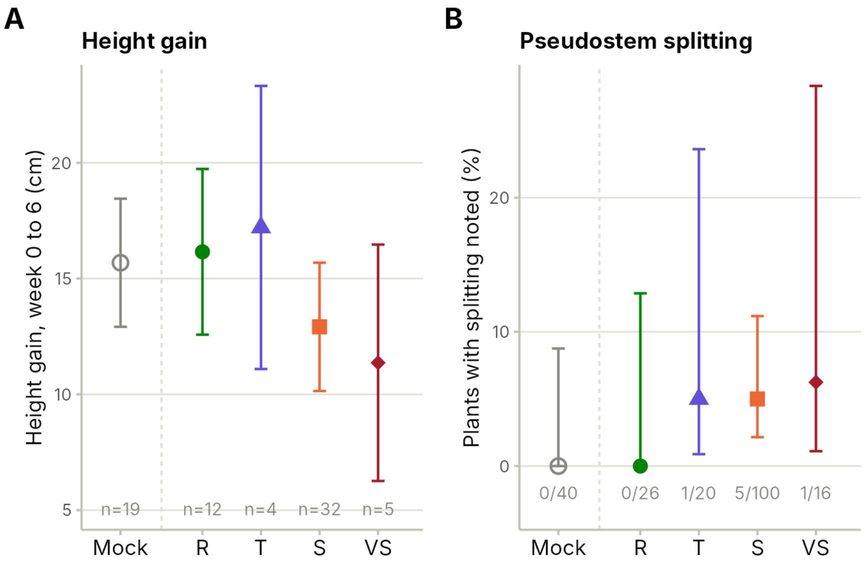
A
Height gain
Height gain, week 0 to 6 (cm)
20
15
10
5
n=19
n=12
n=4
n=32
n=5
Mock
R
T
S
VS
B
Pseudostem splitting
Plants with splitting noted (%)
20
10
0
0/40
0/26
1/20
5/100
1/16
Mock
R
T
S
VS
