## Supplementary figures and images for "Evaluating novel sources of resistance to *Fusarium oxysporum* f. sp. *Cubense* tropical race 4 (TR4) in a global banana diversity panel"

### Supplementary figure S2

Suppl Figure S2

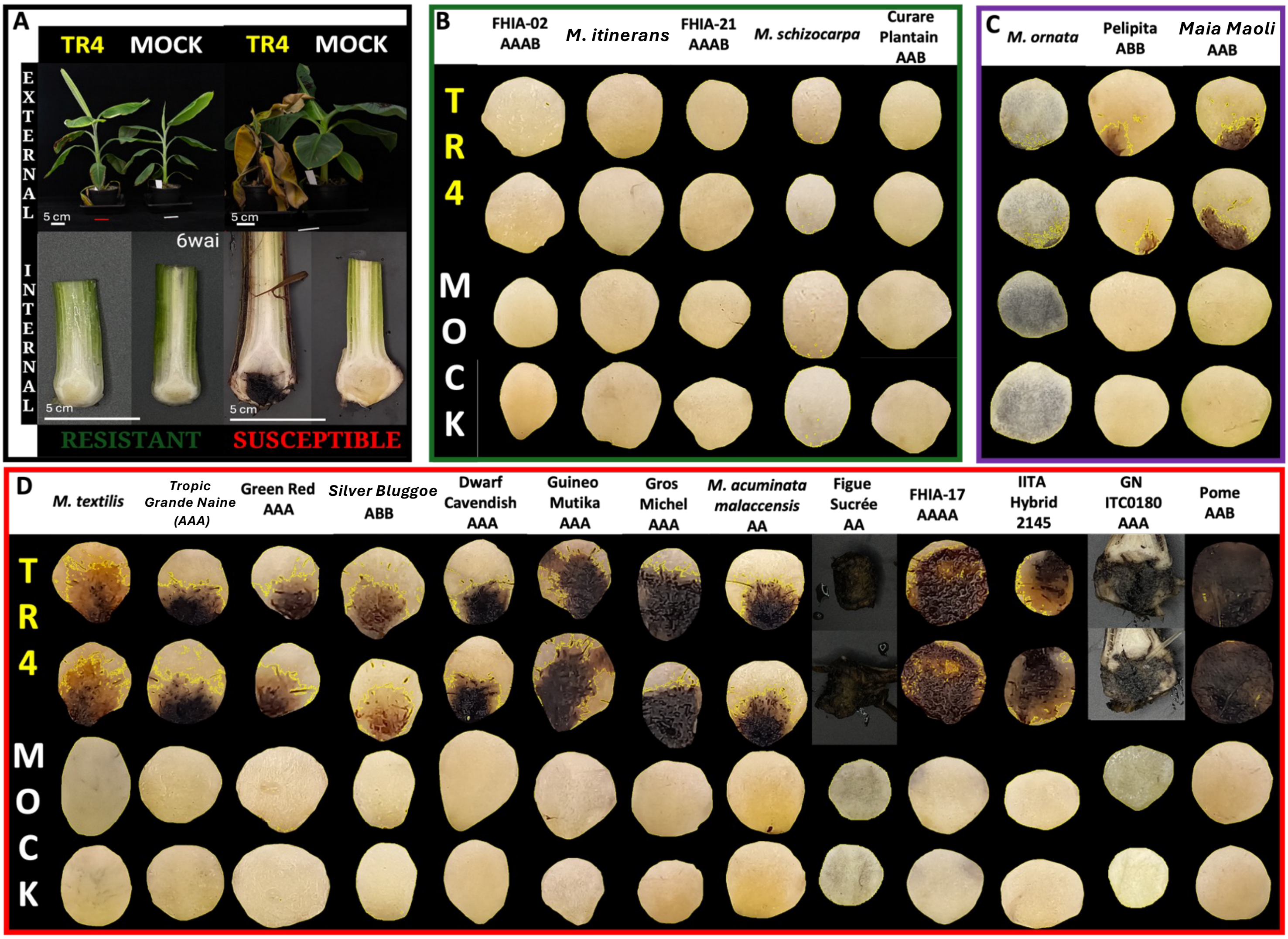

### Supplementary figure S3

#### Suppl Figure S3

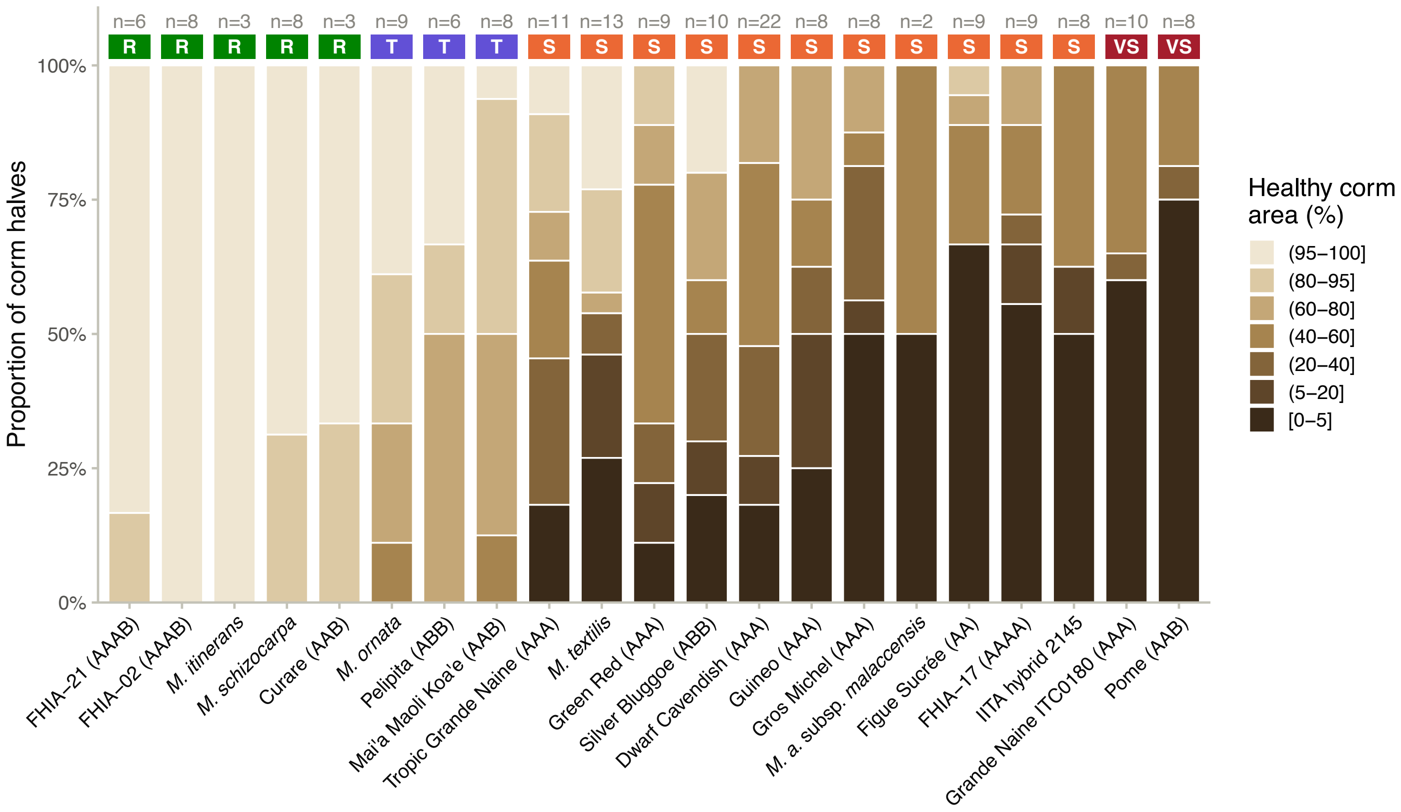

### Supplementary figure S4

Suppl Figure S4

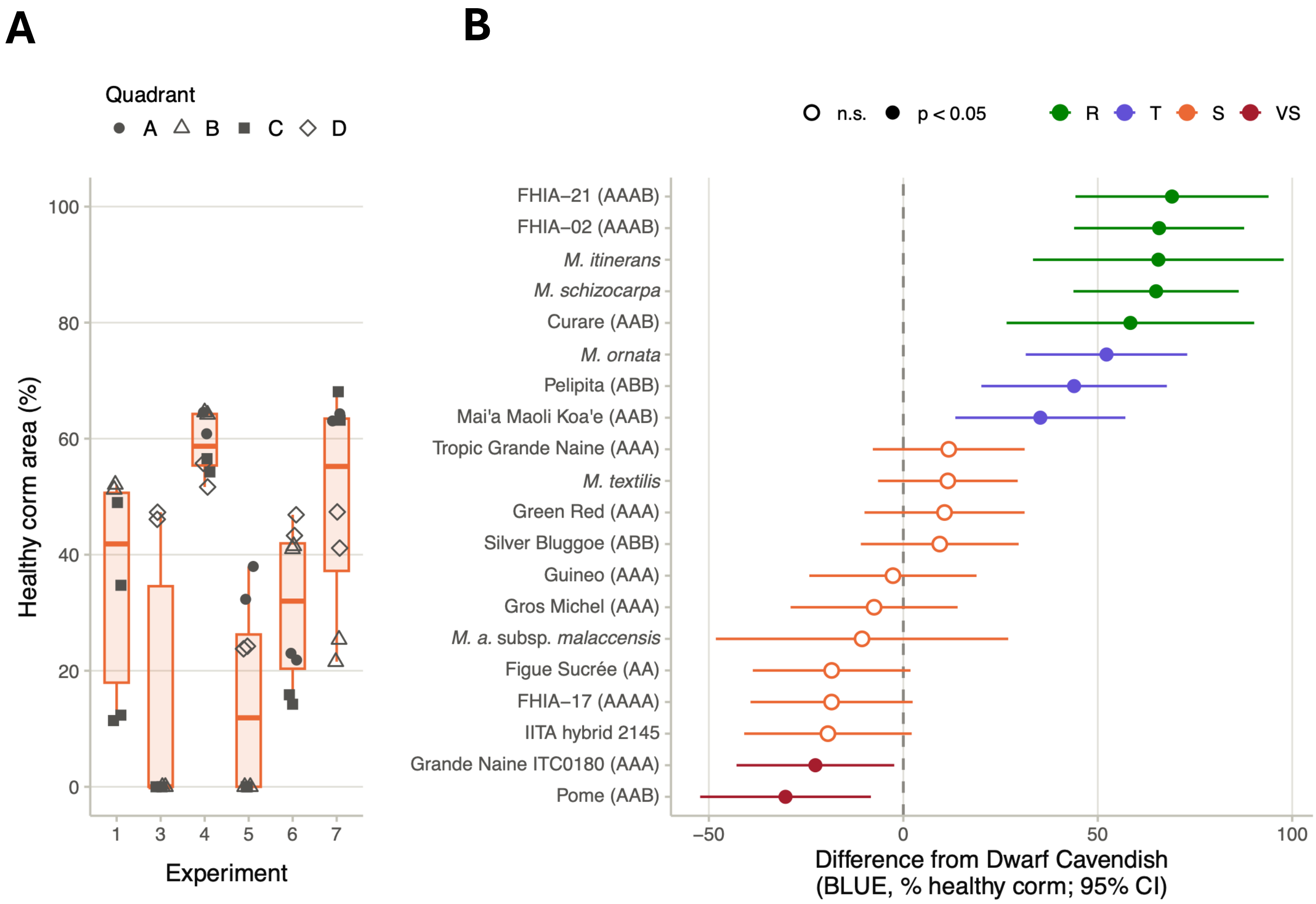

### Supplementary figure S5

<u>Suppl Figure S5</u>

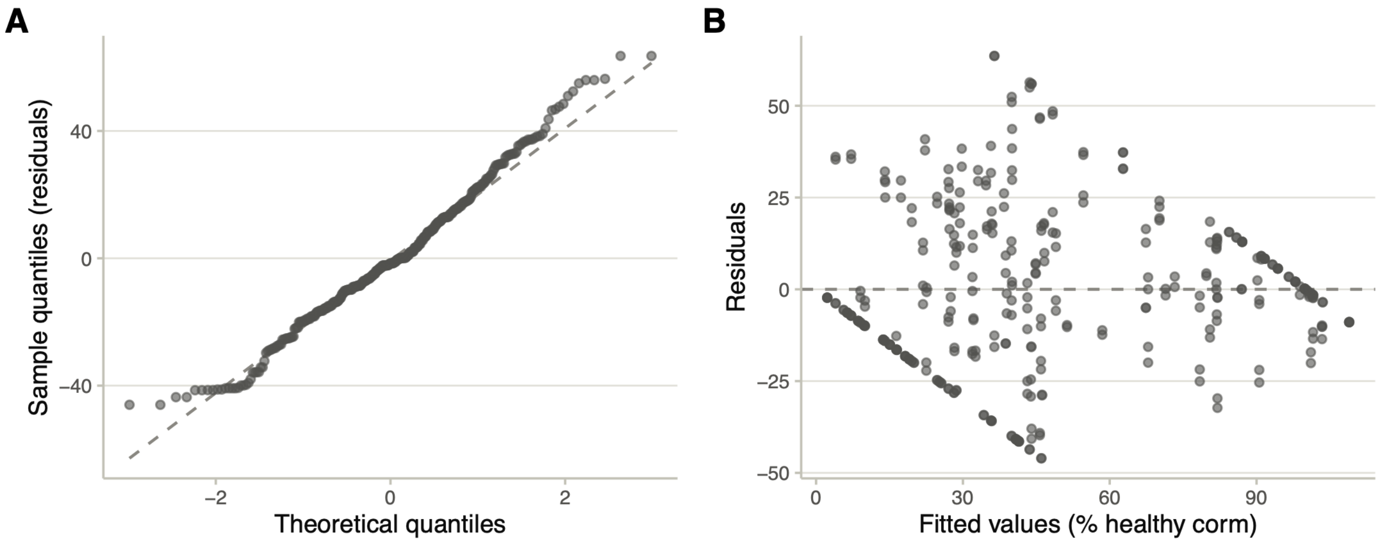
